# Endogenous oncogenic KRAS expression increases cell proliferation and motility in near-diploid hTERT RPE-1 cells

**DOI:** 10.1101/2023.09.08.556827

**Authors:** Naushin L. Hindul, Lauren R. Abbott, Sumaya M.D. Adan, Kornelis R. Straatman, Andrew M. Fry, Kouji Hirota, Kayoko Tanaka

## Abstract

About 18% of all human cancers carry a mutation in the *KRAS* gene making it among the most sought-after anti-cancer targets. However, mutant KRas protein has proved remarkably undruggable. The recent approval of the first generation of RAS inhibitors therefore marks a seminal milestone in the history of cancer research. Inevitably though, it also raises the predictable challenges of limited drug efficacies and acquired resistance. Hence, new approaches that improve our understanding of the tumorigenic mechanisms of oncogenic RAS within more physiological settings continue to be essential. Here, we have employed the near-diploid human hTERT RPE-1 cells to generate isogenic cell lines in which one of the endogenous *KRAS* alleles carries an oncogenic *KRAS* mutation at glycine 12. Cells with a *KRAS^G12V/+^*, *KRAS^G12C/+^*, or *KRAS^G12D/+^* genotype, together with wild-type *KRAS^G12G(WT)/+^* cells, reveal that oncogenic *KRAS.G12X* mutations increase cell proliferation rate, while further analyses showed that *KRAS^G12V/+^* cells had increased cell motility and reduced focal adhesions. EGF-induced ERK phosphorylation was marginally increased in *KRAS^G12V/+^*cells, while EGF-induced AKT phosphorylation was comparable between *KRAS^G12V/+^*and *KRAS^G12G(WT)/+^* cells. Interestingly, the *KRAS^G12V/+^*cells were more sensitive to hydroxyurea and a MEK inhibitor, U0126, but more resistant to a PI3K inhibitor, PIK-90, than the *KRAS^G12G(WT)/+^*cells. A combination of low doses of hydroxyurea and U0126 showed an additive inhibition on growth rate that was greater in *KRAS^G12V/+^*than wild-type cells. Collectively, these cell lines will be a valuable resource for studying oncogenic RAS signalling and developing effective anti-KRAS reagents with minimum cytotoxicity on wild-type cells.

## Introduction

The RAS family of small GTPases acts as a signalling hub, regulating fundamental biological activities through multiple downstream pathways (1). Human *RAS* genes are frequently mutated in cancers, and the germline mutations cause disorders termed RASopathies, underlining their physiological importance (2,3). Among the three human *RAS* genes, *KRAS*, *NRAS* and *HRAS*, the *KRAS* gene exhibits the highest mutation rate, reaching about 18% across all human cancers (Catalogue Of Somatic Mutations In Cancer (COSMIC), v98). Over 80% of the *KRAS* oncogenic mutations are missense mutations at glycine position 12, which is most often mutated to aspartic acid (G12D), valine (G12V) or cysteine (G12C) (4).

Once described as “undruggable,” the oncogenic KRas protein can today be targeted on account of the extensive efforts inspired by the first KRas.G12C inhibitor developed by Shokat and colleagues (5). The KRas.G12C inhibitors are now available for use in the clinic (6,7), while promising KRas.G12D, KRas.G12V and pan-Ras inhibitors have also been developed (8–11). However, there remain significant pending challenges, not least acquired resistance to these inhibitors (12,13). To design rational and effective counteracting strategies, it is crucial to obtain deeper insights on how oncogenic KRas signalling influences cell processes.

Early studies that relied on *RAS* overexpression in cultured cells revealed the high potency of oncogenic *RAS* proteins to cause various cancer-relevant phenotypes, including hyper-activation of downstream signalling components (such as ERK and AKT), cell transformation, cell migration and senescence (14). However, through the elegant analysis of *KRAS* mouse models, the importance of looking at endogenous *KRAS* oncogenic mutations has become well-recognised (14–18). A simpler and cost-effective alternative to mouse models may be examining the consequences of expression of endogenous *KRAS* mutations in human cultured cells derived from normal tissues and immortalised through the expression of human telomere reverse transcriptase (hTERT). This would allow phenotypic changes to be determined in cells that retain a stable and near-diploid karyotype (19,20). Previously, hTERT-immortalised HME1 (mammary epithelial) cells expressing a *KRAS.G13D* mutant and hTERT-IMEC (mammary epithelial) cells expressing a *KRAS.G12V* mutant were generated using adeno-associated viral (AAV) mediated gene targeting (21,22). These studies established that these endogenous oncogenic *KRAS* mutations do not cause cellular transformation, unlike their overexpression (21,22). The studies also reported that the cell proliferation rate (21) or morphology (22) of the cells carrying these endogenous oncogenic *KRAS* mutations were comparable to the parental cells.

Here we report the generation of isogenic human cell lines carrying heterozygous oncogenic *KRAS* mutations using hTERT-immortalised RPE-1 (retinal pigment epithelial cells). However, as the original hTERT RPE-1 cell line (CRL4000^TM^, ATCC) turned out to carry the c.30_35dup *KRAS* mutation at one of the *KRAS* loci, we targeted this locus so that it was converted into the oncogenic *KRAS.G12V*, *KRAS.G12D* or *KRAS.G12C* mutations, as well as generating a wild-type *KRAS.G12G.* These will be referred to here as *KRAS^G12V/+^*, *KRAS^G12D/+^*, *KRAS^G12C/+^* and *KRAS^G12G(WT)/+^*cell lines.

Characterization of these isogenic cell lines revealed that all three endogenous oncogenic *G12X* mutations resulted in increased cell proliferation compared to the G12G(WT) counterpart. Further analyses of the *KRAS^G12V/+^*cell line showed an increase in cell motility and a marginal increase in the EGF-stimulated ERK activation profile. Interestingly, compared to the *KRAS^G12G(WT)/+^* cells, the *KRAS^G12V/+^* cells were more sensitive to hydroxyurea (HU) and a MEK inhibitor, U0126, but more resistant to a PI3K inhibitor, PIK-90. The results highlight the impact of a single endogenous oncogenic *KRAS* mutation and showcase the value of these cell lines as a powerful tool in the study of oncogenic *KRAS*-induced pathogenesis.

## Results

### Identification of a heterozygous c30_35dup mutation in *KRAS* in the hTERT RPE-1 cell line

During the process of establishing a protocol to genotype exon 2 of the *KRAS* gene locus, we found that one of the *KRAS* alleles has the insertion mutation, termed c.30_35dup. As our lab stock of hTERT RPE-1 cells had been obtained more than 10 years ago, we purchased a fresh aliquot of hTERT RPE-1 cells (CRL-4000®) from ATCC in January 2021. However, we found that this too contained the heterozygous c.30_35dup mutation (Fig. 1). The presence of this mutation in these cells has been previously reported (21,23), indicating that the original hTERT RPE-1 cell line likely carries the c.30_35dup mutation in one of the *KRAS* alleles. The c.30_35dup mutation causes the duplication of Ala11_Gly12 (p.Ala11_Gly12dup mutation), resulting in the insertion of two additional amino acids, Ala-Gly, at positions 13 and 14 (Fig. 1B). Considering that Gly12 and Gly13 are hotspots for the *KRAS* oncogenic mutation, the c.30_35dup mutation is highly likely to affect KRas function. In fact, in the Catalogue Of Somatic Mutations In Cancer (COSMIC v.98), 6 cases of the c.30_35dup mutation are reported out of 49887 cases of *KRAS* mutation. Furthermore, enrichment of the active GTP-bound form of KRas in lysates of the hTERT RPE-1 cell line was reported (21). Therefore, in this study, we decided to target the chromosome carrying the c.30_35 *KRAS* mutation to generate isogenic cell lines of *KRAS^G12G^ ^(WT)/+^*, *KRAS^G12V/+^*, *KRAS^G12C/+^* and *KRAS^G12D/+^*.

**Figure 1.**
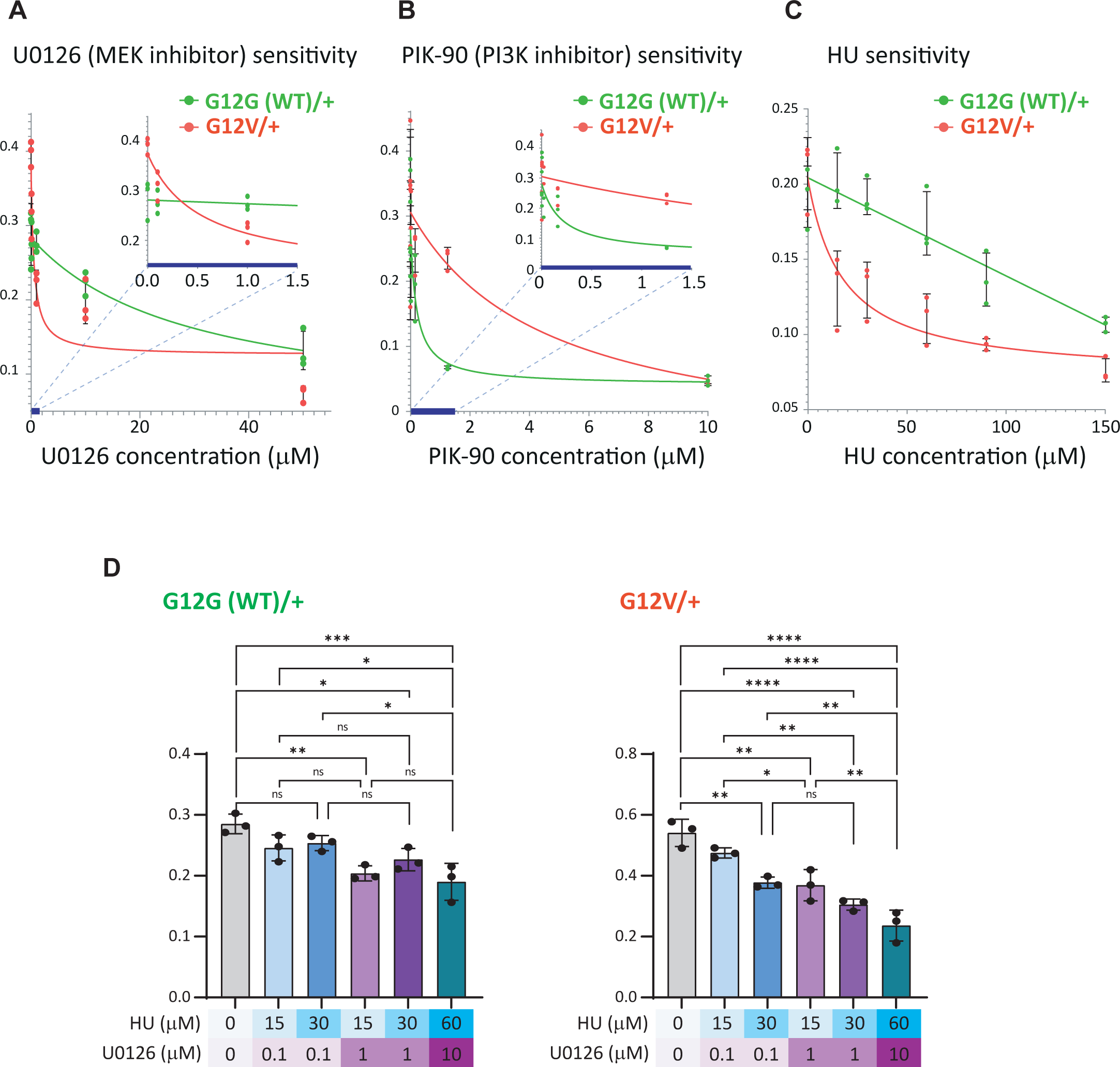
hTERT RPE-1 cells carry a heterozygous c.30_35dup mutation at the *KRAS* gene allele. (A) The PCR-based genotyping of the *KRAS* exon2. Using a pair of primers, AlugenomecheckF and Kras_R, the *KRAS* exon 2 of both of the chromosomes of hTERT RPE-1 cells were amplified. The location of the c.30_35dup insertion mutation is indicated as an orange star in exoon2. The agarose gel image shows successful amplification of the expected gene fragment, indicated by an arrow. (B) The PCR product shown in (A) was sequenced using the primer AlugenomechekF to reveal that one of the chromosomes carries the c.30_35dup insertion mutation. The six nucleotides corresponding to the c.30_35 are orange-underlined, and the inserted duplicated AGCTGG nucleotides are highlighted in orange. The encoded amino acids of the presented region are shown in grey letters below the nucleotide sequences. The c.30_35dup mutation causes the duplication of Ala11_Gly12, leading to the additional Ala_Gly at positions 13 and 14 (shown in orange letters). The Gly12 and Gly13 oncogenic hotspot amino acids are shown in magenta.

### Integration of *KRAS^G12G/C/D/V^* at the endogenous *KRAS* locus in hTERT RPE1 cells

The *KRAS* gene locus was edited through CRISPR-Cas9 mediated recombination of a DNA fragment spanning the *KRAS* exon 2 that encodes Gly12 (Fig. 2A). We used the ER^T2^-Cre-ER^T2^ hTERT RPE-1 cells that we had previously generated (24) to facilitate the removal of the selection marker (histidinol dehydrogenase, hisD), which is sandwiched between two LoxP sites (Fig. 2A, LoxP-hisD-LoxP, or LHL). In this cell line, the expression and activation of the ER^T2^-Cre-ER^T2^ can be induced by the addition of doxycycline and 4-hydroxy tamoxifen (4-OHT) to the culture media, leading to removal of the LHL cassette. The *KRAS* allele was targeted by a CRISPR-Cas9 construct, pX330-kras_CRISPR, and the c.30_35dup mutation was replaced with the G12V, G12D, G12C or G12G (WT) alleles using the pKH-His-DA-Ap plasmid, as previously described (24). We originally intended to generate KRas.G12X inducible cell lines where the presence of the LHL cassette represses the expression of the KRas.G12X. However, when the KRas.G12V expression was examined using an anti-G12V specific antibody, the LHL cassette turned out not to repress the KRas.G12V expression. Cells harbouring the *KRAS.LHL* allele, retaining the LHL cassette, were found to express a similar level of the G12V variant compared to cells harbouring the *KRAS.LoxP* allele from which the LHL cassette had been removed (Supporting information S1). Therefore, we decided to analyse the phenotypes resulting from constitutive expression of endogenous oncogenic *KRAS* mutant, and removed the LHL cassette so that the genomic sequence of the two *KRAS* loci were more comparable (Fig. 2A, B).

**Figure 2.**
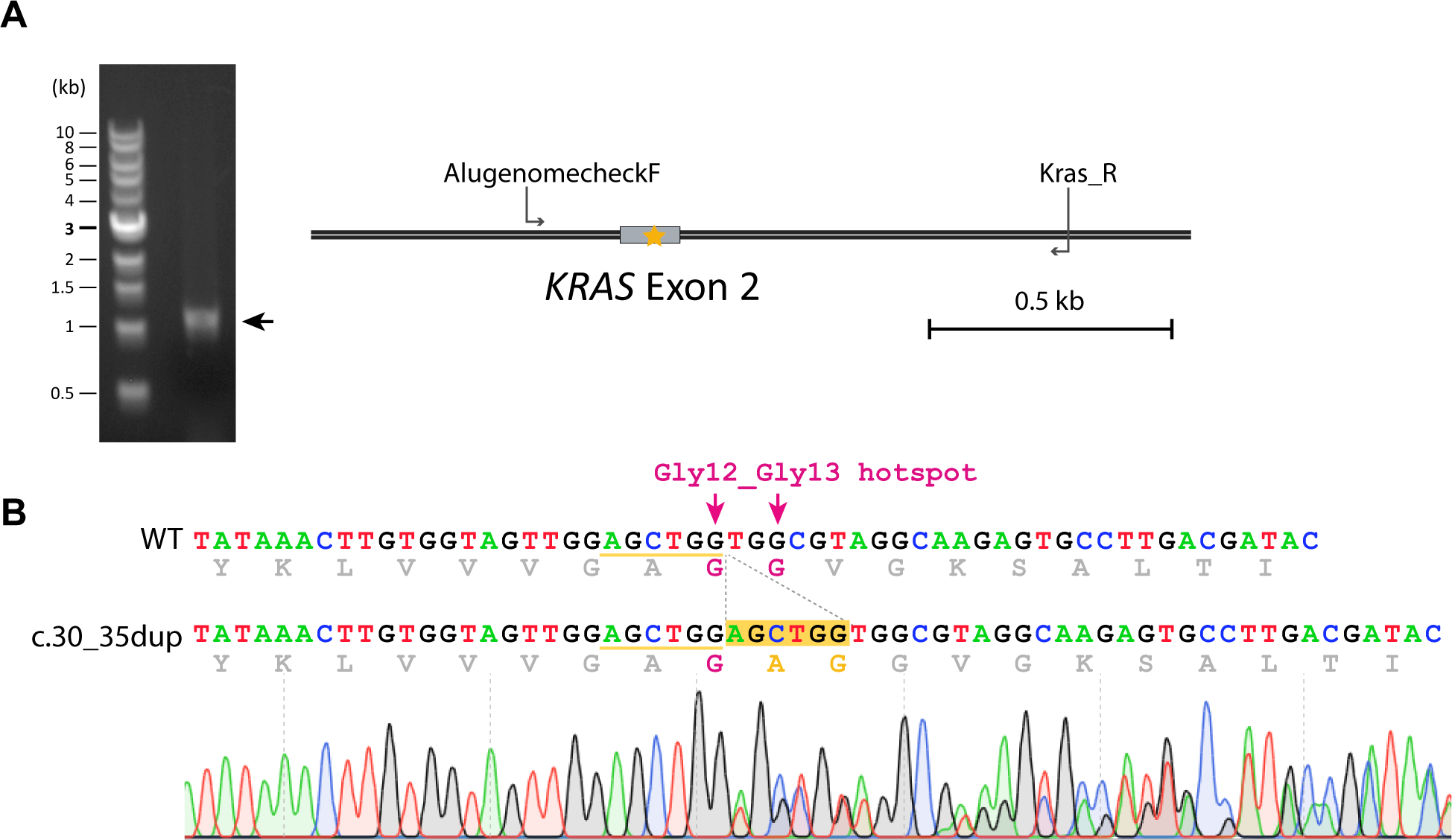
Successful removal of the *KRAS.c.30_35dup* allele to generate isogenic *KRAS^G12X/+^* cell lines. (A) A schematic diagram of chromosome editing of the *KRAS* gene locus. The original unedited *KRAS* locus is shown as *KRAS.unedited*. The parental cell line, hTERT RPE-1, carries one wildtype *KRAS* and one *c.30_35dup* mutant *KRAS* allele. A DNA fragment encoding histidinol dehydrogenase (hisD), flanked by LoxP sites (LoxP-HisD-LoxP, LHL) together with the homologous arms (highlighted in green) was inserted into intron 2 of the *KRAS* gene to create the *kras.LHL* locus. During this process, the c.30_35dup allele can be removed and replaced with the intended *KRAS.G12X* alleles. The generated cells were further treated with doxycycline and 4-OHT so that the LHL cassette was excised, generating the *KRAS.LoxP* allele. A pair of primers, KRAS_intron1_F-2022 and KRAS_R_Nov21, are indicated in the diagram. (B) PCR-based genotyping. The PCR reaction was conducted using primers KRAS_intron1_F-2022 and KRAS_R_Nov21. The primer KRAS_intron1_F-2022 is outside of the fragment used for *KRAS* gene editing. The PCR reaction produced the expected products of 2163 bp for *kras.unedited* and 1908 bp for *kras.LoxP*. For the *KRAS^G12C/+^* cells, both chromosomes carry the *KRAS.LoxP* allele, and the PCR product shows only one band of 1908 bp. However, only one of them has the *KRAS.G12C* mutation, as shown in (C). (C) Sequencing of the genome PCR products confirms the successful introduction of the intended G12X mutations. The PCR products presented in (B) were sequenced to confirm the successful replacement of the c.30_35dup mutation with the intended G12X alleles. Below the nucleotide sequences, corresponding amino acid sequences are shown in grey. The Gly12-Gly13 oncogenic mutation hotspot is indicated in magenta, and the position 12 amino acids are indicated in purple.

We successfully generated one clone of G12G (WT), four clones of G12V, one clone of G12C and one clone of G12D that carry the desired mutations at exon 2 in a heterozygous manner (Fig. 2B). In the following analyses, one of the four G12V clones (clone 19-16) and each clone of the other genotypes were analysed. Sequencing confirmed that the *KRAS* coding sequences (CDSs) of these clones did not contain additional mutations (Supporting information S2A-D). For amplification of the *KRAS* CDS for sequencing analysis, we used a pair of primers that can amplify both *KRAS4A* and *KRAS4B*, two *KRAS* splice variants. In all cell lines, the sequence signals were dominated by the *KRAS4B* transcript, and the *KRAS4A* transcript could not be detected (Supporting information S2A-D). For each mutant cell line, both wild-type and the G12X variant sequencing signals were detected at similar intensities, confirming that all cell lines carry heterozygous *KRAS.G12X* mutations (Supporting information S2A-D).

### KRAS-G12X oncogenic mutations cause an increased rate of proliferation

One of the critical features of cancer cells is the loss of cell cycle control that leads to hyperplasia. All the generated oncogenic *KRAS^G12X/+^*mutant cells showed an increased rate of cell proliferation with the doubling time reduced approximately two-fold compared to the control *KRAS^G12G(WT)/+^* cells (Fig. 3A, B). This demonstrates that one copy of a *KRAS.G12X* oncogenic mutation is sufficient to cause a key cancer-related phenotype. Flow cytometry analysis of the *KRAS^G12V/+^* and *KRAS^G12G(WT)/+^*cells showed that there was an increase in the proportion of cells in S and G2 phase with a concomitant reduction of cells in G1 in the *KRAS^G12V/+^*cell line (Fig. 3C, Supporting information S3). Considering the shorter doubling time of the KRAS^G12V/+^ cells, the time spent in S and G2 phases are estimated to be comparable between the *KRAS^G12V/+^* and *KRAS^G12G(WT)/+^* cells, but the duration of the G1 phase of the cell cycle is substantially reduced in the *KRAS^G12V/+^* cells suggesting loss of restriction point control. Meanwhile, sub-G1 populations in these two cell lines were comparable, indicating that the *KRAS^G12V/+^* mutation does not cause substantial cell death (Fig. 3D, Supporting information S3).

**Figure 3.**
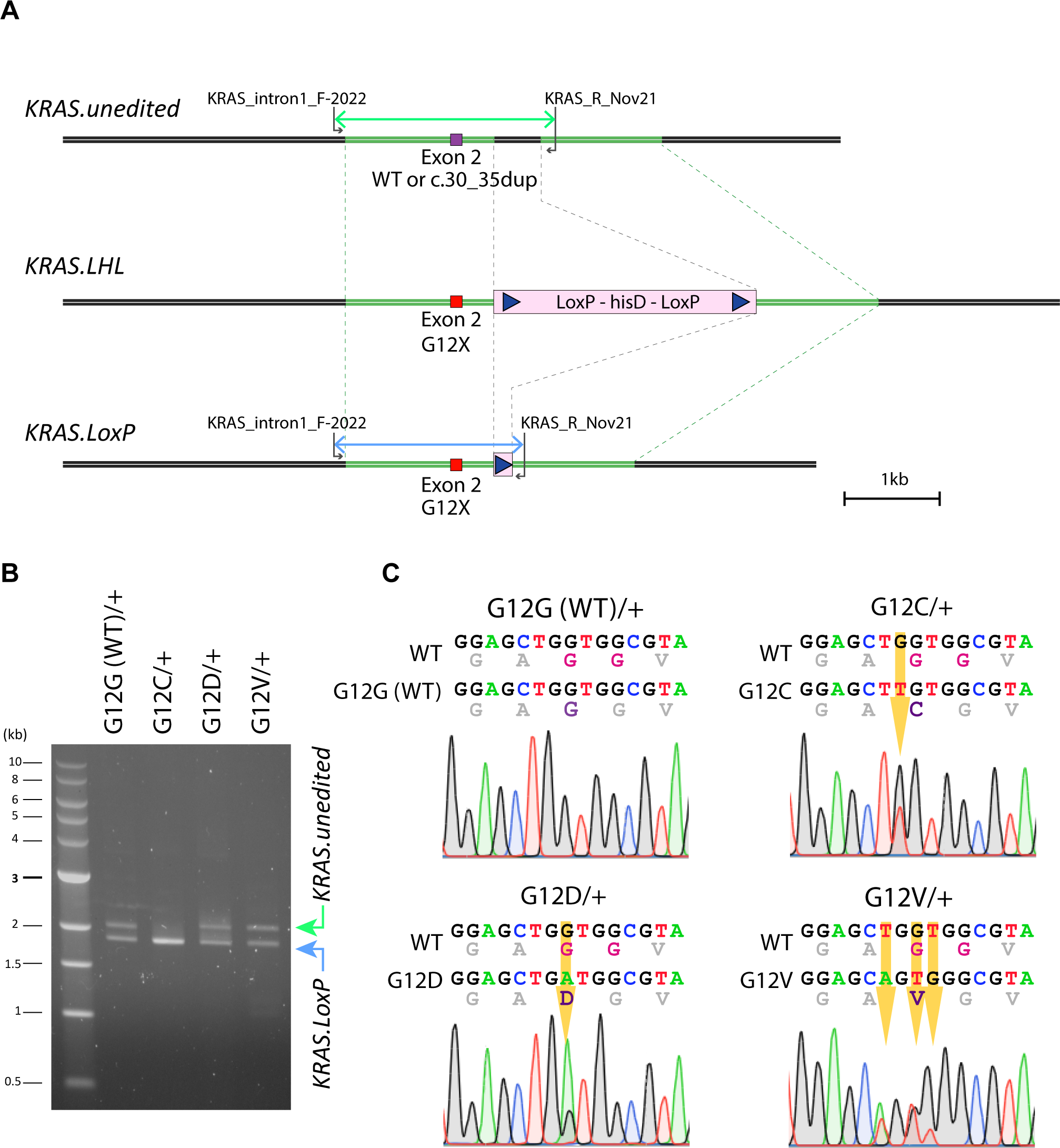
One copy of the endogenous oncogenic *KRAS* mutation causes an increase in cell proliferation. (A) Proliferation rates of the *KRAS^G12X/+^* cells. Exponential growth curves were generated by fitting to three biological replicates using Prism (GraphPad). Y-axis is logarithmic. (B) Oncogenic *KRAS.G12X* mutations substantially reduce the doubling time. From the data presented in (A), the doubling time for each cell line was estimated using Prism (GraphPad). Error bars represent a 95% confidence interval. (C) Distinct cell cycle profiles between the *KRAS^G12V/+^* and the *KRAS^G12G(WT)/+^* cells. Flow cytometry analysis for the DNA contents was conducted for the *KRAS^G12V/+^* and *KRAS^G12G(WT)/+^* cells. A summary result of the biological triplicates presented in Supporting information S3 is presented to show the mean and SD values of the cell cycle populations for each sample. Considering the reduced doubling time of the *KRAS^G12V/+^* cells, as shown in (B), the G1 phase of the cell cycle is estimated to be substantially reduced in the *KRAS^G12V/+^* cells. (D) The sub-G1 population is comparable between the *KRAS^G12V/+^* and the *KRAS^G12G(WT)/+^*cells. The sub-G1 population of the flow cytometry data presented in Supporting information S3 is summarised, and the mean and SD values are presented. Unpaired t-test showed that the two samples showed non-significant differences.

### *KRAS^G12V/+^* oncogenic mutation causes increased cell motility and reduced paxillin structures

Another key phenotype of cancer cells is an increase in migration that contributes to their metastatic potential. The generation of a human HEK293 stable cell line with the expression of an *HRAS.G12V* oncogenic protein suggested that this mutation can cause an increase in cell motility (25). To examine whether the endogenous *KRAS^G12V/+^* oncogenic mutation affects cell motility, live cell imaging was conducted using a label-free ptychographic imaging system. This allowed us to plot cell movement tracks over time revealing substantially enhanced motility for the *KRAS^G12V/+^* compared to *KRAS^G12G(WT)/+^* cells (Fig. 4A). Quantification confirmed the increased velocity of cells carrying the oncogenic *KRAS^G12V/+^* mutation; the mean values are 10.76 nm/sec and 5.20 nm/sec for the *KRAS^G12V/+^*and *KRAS^G12G(WT)/+^* cells, respectively (Fig. 4B). In support of this increased motility phenotype, immunofluorescence microscopy analysis indicated a significant reduction in structures that stained positive for paxillin in the *KRAS^G12V/+^* cells (Fig. 4C). As paxillin is a major component of the focal adhesion complex, these data demonstrate that one copy of the endogenous *KRAS.G12V* oncogenic mutation is sufficient to enhance cell motility and potentially alter cell adhesion properties.

**Figure 4.**
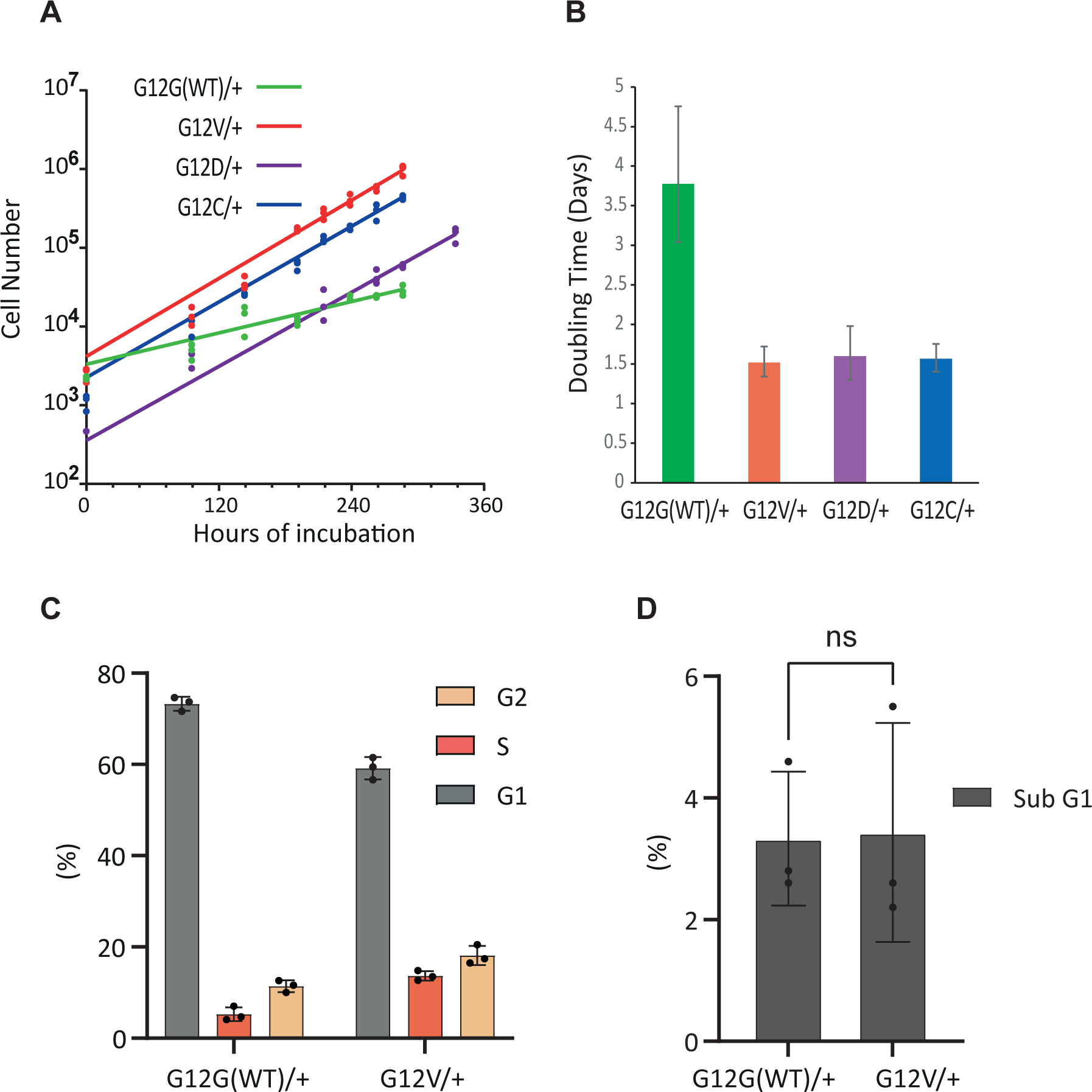
The *KRAS^G12V/+^* cells show enhanced motility with reduced paxillin structures. (A) The *KRAS^G12V/+^* and the *KRAS^G12G(WT)/+^* cells were cultured on collagen-coated plates, and the cell movement was recorded every 20 min for 31 time-frames using ptychographic phase imaging. Each cell movement was tracked and analysed using the Livecyte system (Phasefocus) as described in the Experimental Procedures. Data from three biological replicates from each cell line is presented. The positions of cell tracks are plotted on an X-Y axis, where the start of each track (time 0) is set to the centre of the plot ((x, y) = (0, 0)). Each track is shown in a different colour. (B) The dataset presented in (A) was analysed to deduce the velocity of each cell from one time-frame to the next at every time point. The obtained values are plotted for each biological replicate and are analysed with a nested t-test (Prism, GraphPad), which shows an increase in the *KRAS^G12V/+^* cells. (C) A decrease in the paxillin signals in the *KRAS^G12V/+^* cells. The status of paxillin structures in the *KRAS^G12V/+^* and the *KRAS^G12G(WT)/+^* cells were visualised by immuno-fluorescence microscopy as described in the Experimental Procedures. The actin stress fibre and the nuclei were also counter-stained. The ratio of the area occupied by the paxillin structures and the cytoplasm was deduced for each image, as stated in Supporting information S4. For each biological replicate, 20 images were analysed, and the data from three biological replicates was analysed by a nested t-test (Prism, GraphPad). The area occupied by the paxillin structures is significantly reduced in the *KRAS^G12V/+^*cells.

### Attenuation of ERK signalling is compromised in *KRAS^G12V/+^* cells

It is well accepted that oncogenic KRAS leads to activation of the ERK and AKT signalling pathways (14). Interestingly, treatment of *KRAS^G12V/+^* cells with epidermal growth factor (EGF) only led to a transient, but not a constitutive, activation of ERK and AKT as determined by their phosphorylation (Fig. 5A & B). However, attenuation of the phospho-ERK signal was significantly reduced in the *KRAS^G12V/+^* cells compared to the *KRAS^G12G(WT)/+^*cells. In contrast, attenuation of the phospho-AKT signal occurred at a comparable rate between these two cell lines.

**Figure 5.**
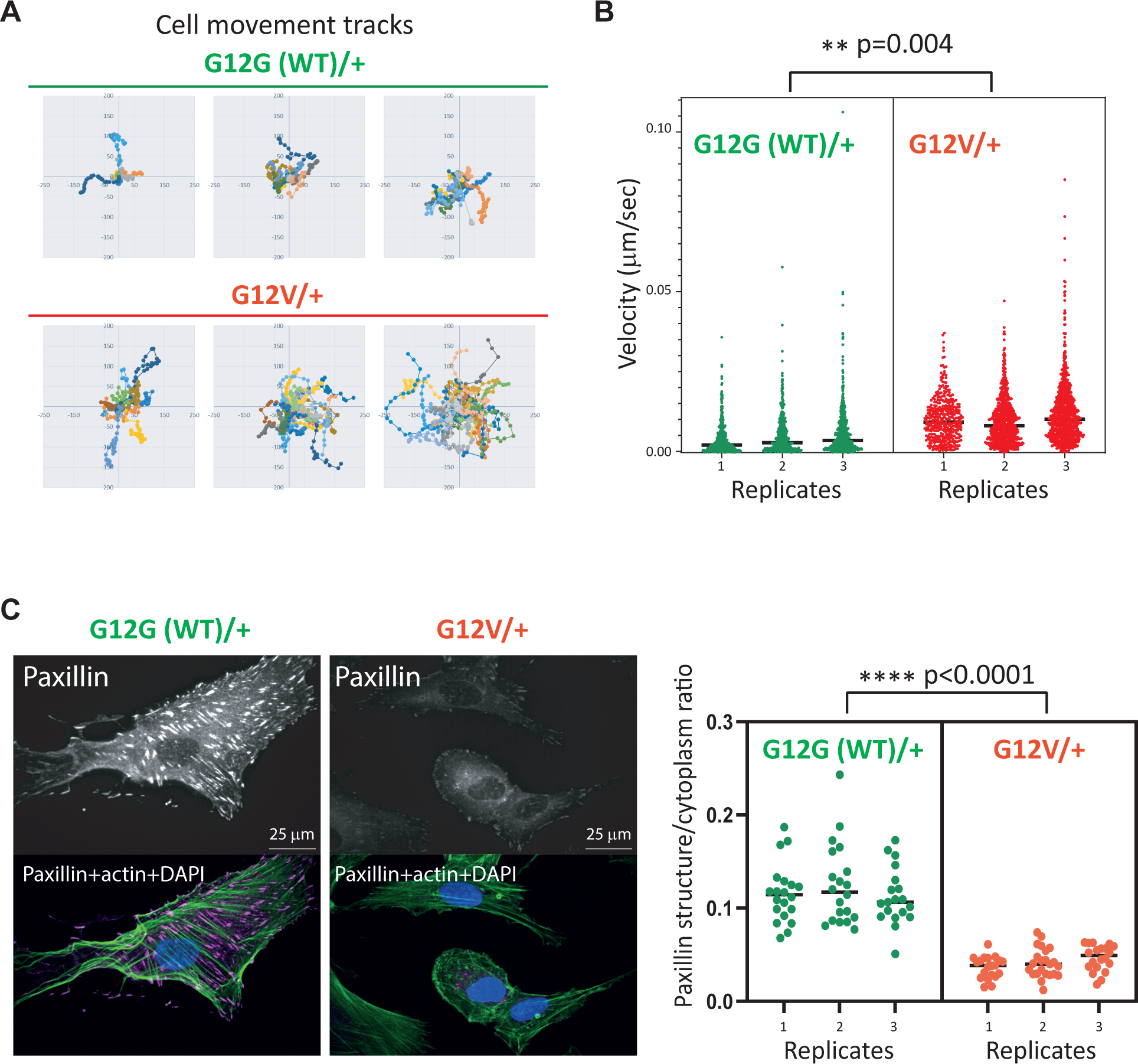
ERK and AKT activation and attenuation profiles upon EGF treatment in the *KRAS^G12V/+^*and *KRAS^G12G(WT)/+^* cells. Cells were serum-starved for 48 hours before the EGF stimulation as described in the Experimental Procedures. ERK (A) and AKT (B) phosphorylation status were monitored for 2.5 min, 5 min, 15 min, 30 min and 45 min after the EGF stimulation by Western blotting. The ratios of phosphorylated ERK1/2 (pERK) and the internal control γ-tubulin (A) or the ratio of phosphorylated AKT (pAKT) and the total AKT (B), were quantitated using the Odyssey imaging system (Li-COR). Three biological replicates, presented in Supporting information S5 and S6, were analysed to plot the graphs that show the mean and the SD values.

### *KRAS^G12V/+^* cells show increased sensitivity to hydroxyurea and a MEK inhibitor but are more resistant to a PI3K inhibitor compared to the *KRAS^G12G(WT)/+^*cells

We next explored whether one copy of the endogenous *KRAS.G12V* mutation might alter drug sensitivity using inhibitors known to target Ras signalling pathways. Interestingly, the *KRAS^G12V/+^* cells exhibited an increase in sensitivity towards U0126, a MEK inhibitor, but were more resistant to PIK-90, a PI3K inhibitor, as compared to the *KRAS^G12G(WT)/+^* cells (Fig. 6A, B). We also examined sensitivity against hydroxyurea (HU), a drug that interferes with DNA replication by inhibiting ribonucleotide reductase. The overexpression of oncogenic *HRAS.G12V* in human cells is reported to cause DNA hyper-replication and hyper-transcription, leading to replication stress (26–28). Hence, we suspected the sensitivity towards HU may be altered in the *KRAS^G12V/+^* cells. Indeed, the *KRAS^G12V/+^*cells showed an increase in sensitivity towards HU, suggesting a defect in DNA replication control in these cells (Fig. 6C). Interestingly, HU and U0126 showed an additive inhibitory effect when applied simultaneously to the *KRAS^G12V/+^*cells (Fig. 6D).

**Figure 6.**
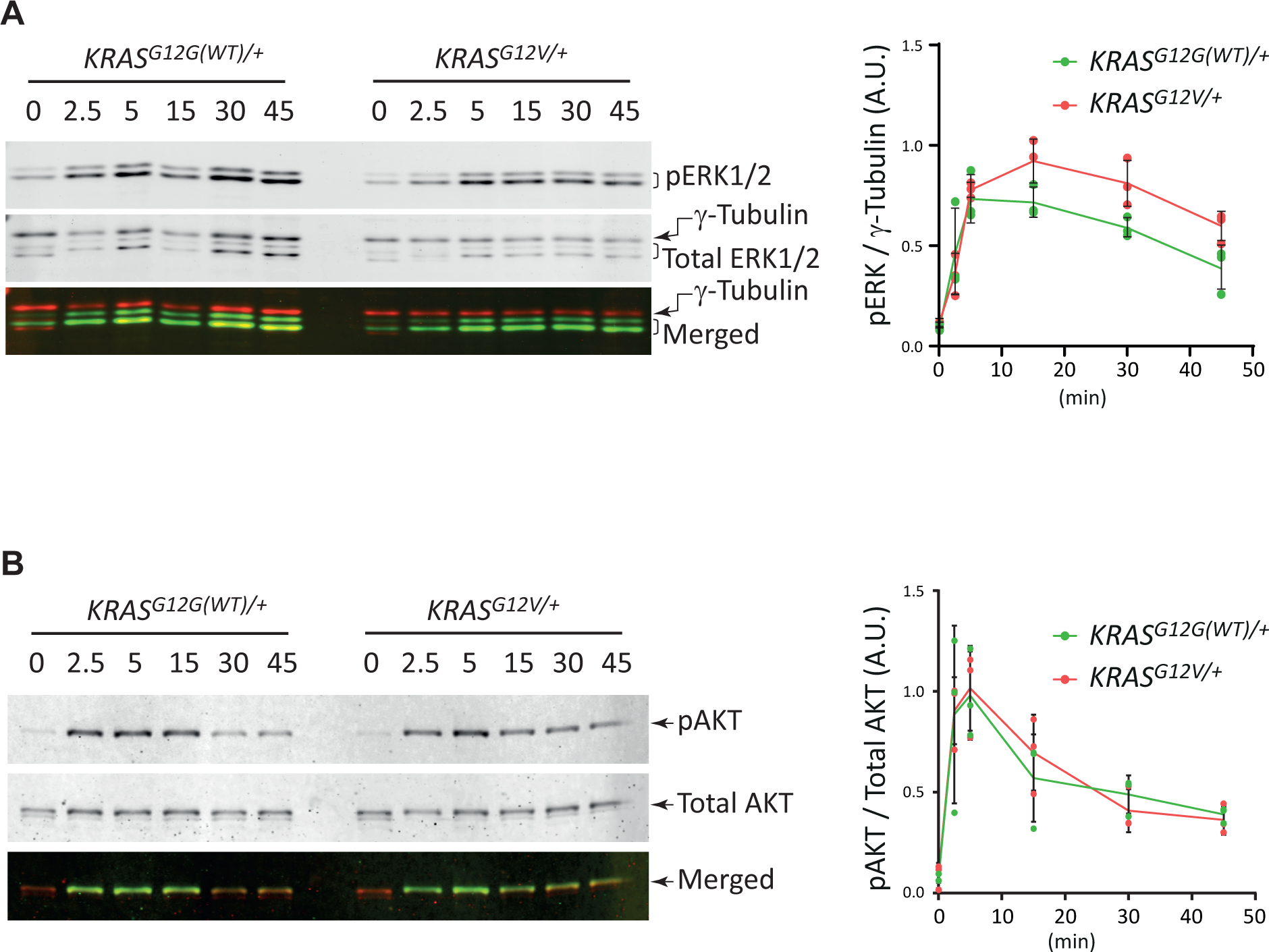
Altered sensitivities of the *KRAS^G12V/+^* and *KRAS^G12G(WT)/+^*cells to U0126, PIK-90 and HU. Crystal violet-based cytotoxicity assays were conducted to monitor the sensitivities towards U0126 (A), PIK-90 (B), HU (C) and the combination of U0126 and HU (D). For (A) – (C), three biological replicates were analysed to generate the graph that shows the mean and SD values, together with the line representing the non-linear fit of data produced by Prism (GraphPad). (A) Cells were exposed to the indicated concentrations of U0126 for 4 days before the fixation, staining and cell lysis. The best-fit IC50 values for the *KRAS^G12V/+^* and *KRAS^G12G(WT)/+^* cells are 0.5268 µM and 28.89 µM, respectively, and the R-squared values were 0.8546 and 0.8789, respectively. (B) Cells were exposed to the indicated concentrations of PIK-90 for 8 days before the fixation, staining and cell lysis. The best-fit IC50 values for the *KRAS^G12V/+^* and *KRAS^G12G(WT)/+^* cells are 4.547 µM and 0.2019 µM, respectively, and the R-squared values were 0.7173 and 0.7870, respectively. (C) Cells were exposed to the indicated concentrations of HU for 7 days before the fixation, staining and cell lysis. The best-fit IC50 value for the *KRAS^G12V/+^* cells was 17.27 µM, whereas the IC50 was calculated “unstable” for the *KRAS^G12G(WT)/+^*cells. The R-squared values were 0.8560 and 0.8131, respectively. (D) The cells were subjected to six different combination treatments of HU and U0126, as shown in the graphs, for 6 days. Data for three biological replicates were obtained and analysed with one-way ANOVA, followed by the Tukey post-hoc test to compare the mean of each case with the mean of every other case. The post-hoc test outcomes are shown in the diagrams except for all the neighbouring cases, which show non-significant differences. All the statistical analyses were conducted using Prism (GraphPad).

## Discussion

*KRAS* is among the most frequently mutated genes in human cancers. For over 40 years, extensive and continuous efforts have been made to understand the disease mechanisms initiated by mutant *KRAS* and to develop interventional therapeutics (29,30). The recent development of RAS inhibitors marks a milestone and promises a brighter future for many cancer patients. It also redefines the current and future challenges, including improving inhibitor efficacies, mitigating acquired resistance, and developing new inhibitors that cover all oncogenic RAS isoforms (11,31,32). To address these issues, obtaining better insights into the mechanisms of oncogenic RAS signalling is essential. The set of isogenic cell lines we developed in this study provides a new human cell culture model with defined genetic elements where the impact of one copy of an endogenous oncogenic *KRAS* mutation can be studied. As the cell lines harbour the inducible ER^T2^-Cre-ER^T2^ cassette, additional conditional gene editing is possible for future analyses (24).

The cell lines carrying the *KRAS.G12X* oncogenic mutations all showed an increase in the rate of cell proliferation compared to the *KRAS.G12G(WT)* control cell line, demonstrating that one copy of the endogenous *KRAS.G12X* oncogenic mutation is sufficient to induce this cancer relevant phenotype in hTERT RPE-1 cells. Interestingly, previous studies that targeted the endogenous *KRAS* allele using an AAV vector to introduce the *KRAS.G13D* mutation in hTERT-HME1 cells (21) and the *KRAS.G12V* mutation into hTERT-IMEC cells (22) reported that the mutations did not increase cell proliferation or changes in cell morphology. This may be due to differences in the host cell lines, and this point will be important to consider when planning future experiments.

Examination of their proliferation rate and cell cycle profile indicated that the G1 phase of the *KRAS^G12V/+^* cells was substantially shorter compared to the *KRAS^G12G(WT)/+^* cells. A similar phenotype was previously observed upon induction of *HRAS.G12V* expression in hTERT RPE1 cells (33). The shortened G1 phase is highly suggestive of a weakened restriction point that could result from an increase in Cyclin D1 expression, which occurs upon oncogenic RAS overexpression (33,34). However, we were not able to detect a significant change in Cyclin D1 expression in these cells (data not shown). Elucidating the mechanism underlying the shortened G1 phase is crucial to understanding how oncogenic *KRAS* increases the proliferation rate even when expressed from the endogenous loci.

An increase in cell motility, another cancer-related phenotype, was also observed in *KRAS^G12V/+^* cells. Interestingly, the cells showed a reduced number of structures that stained for paxillin, suggesting that oncogenic KRAS signalling negatively impacts the focal adhesion complex assembly or maintenance, leading to increased cell motility. This observation is consistent with a previous study where deletion of the *KRAS.G13D* mutation from the endogenous locus of the human colorectal adenocarcinoma cell line, HCT116, results in an increase in paxillin structures (35). A reduction in paxillin structures can also be achieved by MEK inhibition in the parental HCT116 cells (35), indicating that the Raf-MEK-ERK signalling axis plays an important role in the oncogenic Ras-induced reduction in paxillin structures. Involvement of the Raf-MEK-ERK signalling axis was also suggested by the observations that sustained Raf activation induces Paxillin phosphorylation at Ser126 and S130 (36), and active ERK is localised to and facilitates disassembly of focal adhesion complexes (37). However, the precise process of how active ERK facilitates focal adhesion complex disassembly still needs to be addressed, and we believe that our newly generated cell lines serve as an ideal experimental model.

Compared to the cell proliferation and motility phenotypes, the effect of one copy of the endogenous *KRAS.G12V* mutation on the ERK and AKT activation and attenuation profiles upon EGF treatment was somewhat milder. ERK activation was marginally increased and prolonged in the *KRAS^G12V/+^* cells compared to the control *KRAS^G12G(WT)/+^* cells, whereas the AKT activation and attenuation profiles of these two cell lines were comparable. These observations contrast with those in systems where oncogenic RAS is overexpressed. However, they are in line with those in mouse embryonic fibroblasts (MEFs) isolated from the *KRAS.G12D* knock-in mouse model, where serum-stimulated MEK and AKT activation-inactivation profiles are similar between *KRAS.G12D* and control MEFs (38). Considering the phenotypes of increased cell proliferation and motility, these results were surprising. However, we have not explored the cellular localisations of ERK and AKT signalling pathway components. It will be important to obtain the spatio-temporal activation status of ERK and AKT to fully understand the effect of endogenous oncogenic KRAS expression.

Having seen the relatively mild effect on ERK and AKT signalling, it was interesting to observe the different sensitivities to the MEK inhibitor, U0126, and the PI3K inhibitor, PIK-90, of the *KRAS^G12V/+^* and *KRAS^G12G(WT)/+^* cells. The *KRAS^G12V/+^* cells were more sensitive to U0126, potentially indicating that the cells were more reliant on (or “addicted to”) the increased ERK signalling pathway. On the other hand, the *KRAS^G12V/+^* cells were more resistant to PIK-90. Although the mechanisms behind these phenotypes are unclear, the results highlight the importance of also considering the cytotoxicity of healthy cells when developing reagents against oncogenic KRAS signalling. The set of cell lines generated in this study can be a powerful tool to screen drug candidates that fulfil both high efficacy towards the oncogenic KRas and minimum cytotoxicity towards healthy cells.

The increased HU sensitivity of the *KRAS^G12V/+^* cells is consistent with the previous observation that oncogenic RAS expression causes replication stress (26,27,33,39). Possible causes of the replication stress may include increased transcriptional activity (27), which may in turn be a consequence of hyper-activation of ERK signalling. Therefore, we examined whether simultaneous treatment with U0126 (MEK inhibitor) mitigates the HU sensitivity. Strikingly, U0126 exhibited an additive effect with HU on inhibition of growth of the *KRAS^G12V/+^* cells, rather than mitigating the HU sensitivity. This finding may open a new approach to specifically target cells harbouring oncogenic *KRAS* mutations.

In summary, we have generated a set of isogenic *KRAS^G12X/+^*cell lines that will be highly valuable for further functional studies of oncogenic KRas. We demonstrated that one copy of the endogenous *KRAS.G12X* oncogenic mutations causes cancer-relevant phenotypes, including an increase in cell proliferation and motility. This set of cell lines can also be a powerful tool to run pharmacological screens for the development of novel KRas inhibitors with minimum cytotoxicity to healthy non-cancerous cells.

## Experimental Procedures

### Plasmid cloning

Gene editing at the *kras* locus was conducted as previously described (24). Two plasmids, pX330-kras_CRISPR and pKH-His-DA-Ap_kras_integration, were used to insert a LoxP cassette within the *kras* gene. The CRISPR plasmid, pX330-kras_CRISPR, is a derivative of pX330 (Addgene #42230) (40) where a *kras-*targeting guide RNA sequence, 5’ GTATTTCAGAGTTTCGTGAG 3’, has been inserted. The pKH-His-DA-Ap_*kras*_integration plasmid contains a DNA fragment encoding histidinol dehydrogenase under a PGK promoter, which is flanked by a pair of LoxP sites. This LoxP cassette is sandwiched by *kras* homologous arms. The left (5’) arm spans position 9447 to 10951 of chromosome 12, carrying an oncogenic G12V mutation in exon 2, and the right (3’) arm spans position 11399 to 12753 of chromosome 12. The pKH-His-DA-Ap_*kras*_integration plasmid was further mutated to carry other oncogenic mutations, G12C and G12D, as well as wildtype G12G.

### Cell culture, transfection and drug treatment

To culture hTERT RPE-1 cells (ATCC, CRL-4000), DMEM/F-12 (Gibco #31331-028) containing 0.5% sodium bicarbonate, supplemented with 10% fetal bovine serum (Gibco #10500-064), 1% (v/v) Penicillin-Streptomycin (Gibco #15140-122) and 10 µg/ml Hygromycin B Gold (InvivoGen, #ant-hg-1) was routinely used. To culture hTERT RPE-1 ER-Cre-ER cells (24), the media was supplemented with puromycin at a final concentration of 7 µg/ml. Cells were cultured at 37°C, 5% CO_2_. Transfection was conducted using Lipofectamine 3000 Reagent (Invitrogen, #L3000001) following the manufacturer’s instruction. To select clones with a LoxP cassette integration at *kras* gene locus, L-histidinol dihydrochloride (Sigma-Aldrich, #H6647) was added to the media at a final concentration of 1.35-1.5 mg/ml. For doxycycline treatment, doxycycline (Alfa Aesar, #J60422) was dissolved in H_2_O at 1 mg/ml and used at a final concentration of 1 μg/ml in the medium. For 4-OHT treatment, 4-OHT (Merck, #H6278) was dissolved in ethanol at 1 mM and used at a final concentration of 0.5 μM in the culturing medium.

For drug treatment experiments, following inhibitors were used: Hydroxyurea (HU Fluorochem #F043351-1G), dissolved in H_2_O at 20mM, used at final concentrations of 0, 15, 30, 60, 90 and 150 μM, MEK inhibitor (U0126 Apollo Scientific #BIU1095), dissolved in DMSO at 100mM, used at final concentrations of 0, 0.1, 1, 10 and 50 μM, and PI3K inhibitor (PIK-90 TOCRIS Bio-Techne #S1187), dissolved in DMSO at 1mM, used at final concentrations of 0, 2.5, 20, 160, 1250 and 10000 nM.

### PCR based genotyping

For genome extraction, about 1 x 10^6^ cells were collected and were lysed in 400 µl of lysis solution (100 mM NaCl, 25 mM EDTA, 20 mM Tris-Cl (pH 8.0), 0.5% SDS, 0.5% 2-Mercaptoethanol, 0.2 mg/ml Proteinase K (Thermo Scientific EO0491)). The sample was incubated at 55°C for 18-24 hours. On the following day, 450 µl of phenol:chloroform:isoamyl alcohol 25:24:1 was added, and the sample was vigorously mixed. It was centrifuged, and then the upper aqueous was transferred to a fresh tube containing 180 µl of 7.5 M ammonium acetate and 750 µl of 100% Ethanol to precipitate the genome. The sample was centrifuged, and the pellet was air-dried and dissolved in 50µl of Tris buffer (Thermo Scientific, #K0721).

To amplify the *KRAS* locus, a pair of primers Kras_intron1_F-2022(5’ AGCCACCGTGCCCGGCTCACTTGC 3’)and KRAS R_Nov22 (5’ GGAGGTCTTTGAGATTAAATAAATCCTCATCTGCTTG 3’) were used. A typical PCR reaction mix was prepared in 25 µl reaction scale by assembling the following components; 1 µl of extracted genome DNA (11-35 ng/µl), 0.75 µl of each primer (10 µM), 0.75 µl of dNTPs (10 mM each), 0.5 µl of KAPA HiFi DNA polymerase (Roche, #KK2101), 5 µl of 5 x KAPA HiFi buffer (Roche, #KK2101) and 16.75 µl H_2_O. The PCR condition was set as follows; 3 min denaturation at 95°C, 35 cycles of “20 sec 98°C, 15 sec 65°C, 90 sec 72°C” and 5 min 72°C. The amplified fragments were sequenced using the primer AlugenomecheckF (5’ TCATTACGATACACGTCTGCAGTCAACTGG 3’).

### Preparation of cDNA

To examine the coding sequence of the edited *KRAS* gene locus and to confirm the heterozygous expression of the wildtype and the edited *KRAS* gene loci, total RNA was isolated using Direct-zol RNA miniprep kit (ZYMO research, #R2051 and #R2050-1-50) following the manufacturer’s instruction. The mRNA quality was examined by Agilent Bioanalyser to confirm the RNA Integrity Number (RIN) to be 9.5-10. About 5 µg of the purified mRNA was used to generate cDNA using the Go script kit (Promega #A5001). The KRAS coding region was amplified using a pair of primers Kras_Exon1-F (5’ CCGCCATTTCGGACTGGGAGCGAGCGC 3’) and Kras_3’UTR_S (5’ CTGCATGCACCAAAAGCCCCAAGACAGAAATCTTAGG 3’) and the sequencing was conducted using the primers Kras_3’UTR_S and AlugenomecheckF.

### Preparation of cell extracts of growth-factor stimulated cells

*KRAS.G12G* and *KRAS.G12V* hTERT RPE-1 cells were grown to 50% confluent in six-well plates (TPP, #92006). The cells were then serum-starved in the DMEM/F-12 media for 48 hours. The starved cells were growth-factor stimulated with fresh DMEM media containing 100ng/ml epidermal growth factor (EGF Human #A63411-500 (Antibodies.com)), and time-course samples were collected at 0, 2.5, 5, 15, 30 and 45 minutes after the EGF stimulation. To collect cell extracts, the media was removed and cells were washed once with 3ml of phosphate-buffered saline (PBS) and lysed in 60 µl of the modified 3x Laemmli sample buffer that contains 8M urea (192 mM Tris-Cl (pH 6.8), 4.8% SDS (w/v), 24% glycerol (v/v), 1.8 M β-mercaptoethanol, 0.0048% Bromophenol Blue, 8M urea). Before loading samples into the 12% SDS-PAGE gel, they were heat-denatured at 95°C for 5 minutes.

### Western blotting

Proteins were resolved by SDS-PAGE and transferred to 0.2 μm nitrocellulose membranes (Bio-Rad, #1620112). The membranes were blocked overnight at room temperature with a blocking buffer (Li-Cor Intercept Blocking Buffer TBS, #927-60001). The following primary antibodies were used: anti-AKT (Cell Signalling Technology, 40D4 Mouse mAb #2920, dilution 1/2000), anti-pAKT (Cell Signalling Technology, rabbit Antibody #9271, dilution 1/1000),anti-ERK (Thermo-Fisher Scientific: Invitrogen, mouse antibody #10221703, dilution 1/500), anti-pERK (Cell Signalling Technology, rabbit mAb #4370, dilution 1/2000), anti-KRas antibody (OriGene, mouse monoclonal #CF801672, dilution 1/1000), anti-Ras.G12V antibody (Cell signalling technology, rabbit monoclonal #14412, dilution 1/800), anti-γ-tubulin antibody (SIGMA-ALDRICH, mouse monoclonal, #T6557, 1/5000 dilution) and anti-Rgl2 antibody (Novus biologicals, mouse monoclonal, #H00005863-M02, dilution 1/500). To detect the primary antibodies, goat anti-mouse IgG antibody (Li-Cor, #926-68020) and IRDye^®^ 800CW goat anti-Rabbit IgG Antibody were used as a secondary antibody. Membrane images were acquired by Odyssey Infrared Imaging System (Li-Cor Biosciences). The protein levels were quantified using Image Studio software (Li-Cor Biosciences). Three biological replicates were conducted and analysed by Prism software (GraphPad).

### Cell proliferation assays

Cell proliferation status was examined using CellDrop (DeNovix) as previously described (24). Briefly, cells were seeded in six-well plates (TPP, #92006), and cell numbers were counted every 24-96 hours for up to the total incubation time of 360 hours by trypsinising cells and staining them with 0.4% trypan blue solution (Sigma-Aldrich T8154).

### Cell growth inhibition assays

Cell growth inhibition assays were conducted essentially following the previously reported protocol (41). Briefly, cells were grown in 6-well plates (Sarstedt, #83.3920) till 60-80% confluency, washed once with PBS, and were stained with crystal violet staining solution (0.5% crystal violet powder (Sigma-Aldrich C6158), 20% methanol) for 10 min at room temperature. The staining solution was discarded, and the cells were gently but thoroughly rinsed with H_2_O. The stained cells were air-dried before they were lysed with the lysis buffer (0.1 M sodium citrate, 50% ethanol, pH 4.2). The samples were incubated for 30 min at room temperature, gently shaking to ensure complete lysis of the cells. The generated samples were examined by a spectrophotometer at 590 nm absorbance. Three biological replicates were analysed by Prism software (GraphPad).

### Analysis of DNA content by flow cytometry

Cells were trypsinised, harvested by gentle centrifugation and washed with PBS. They were fixed by re-suspending in 70% cold ethanol, and the samples were stored at −20°C until the measurement. The stored samples were washed with PBS three times and were incubated in the staining solution (PBS containing 5mM of EDTA, 20 µg/ml of propidium iodide (Sigma-Aldrich #81845) and 200 µg/ml RNAse A (New England Biolabs #T3018-1)) for four hours on ice. The resultant samples were measured with FACSCanto II (BD Biosciences), and the data analysis was done using BD FACSDiva (BD Biosciences).

### Quantitative live cell imaging

The modes of cell migration were quantitatively measured using the Livecyte kinetic cytometer 2 (Phasefocus) which allows label-free imaging. Cells in 1 ml media were seeded in a 24-well plate coated with Collagen Type 0 (Jellagen, #Jel1018) 48 hours before imaging. Cell images were taken every 20 minutes intervals using a PLN 10 x / NA 0.25 objective for 10 hours at 37°C in an atmosphere supplemented with 5% CO_2_. Acquired images were analysed for instantaneous velocity, migration speed, and cell displacement using Livecyte Analyse software (Phasefocus).

### Immunofluorescence microscopy

Cells were cultured on collagen-coated coverslips (BioCoat® Collagen I 22 mm Round #1, Falcon #CF905) and prior to the fixation, cells were rinsed with PBS. The cells were fixed with freshly prepared 3.7% paraformaldehyde for 5 minutes at room temperature. The cells were then incubated in 0.2% Triton X-100 for 2 minutes to increase cell permeability and were quenched with 1mg/ml NaBH_4_ for 4 minutes, followed by 1 hour of incubation at room temperature in 0.1M Glycine solution. The prepared samples were blocked in the blocking medium (PBS supplemented with 4% fetal bovine serum (Gibco #10500-064) and 1% bovine serum albumin (BSA)(Fisher Scientific #BPE9702-100)) overnight. The samples were then incubated with appropriate primary antibodies diluted in PBS supplemented with 1% (w/v) BSA for 1 hour at 37°C. The following primary antibodies were used in this study; anti-paxillin antibody (BD Biosciences #610052, mouse monoclonal, dilution 1/500) and anti-Rgl2 antibody (Novus biologicals, mouse monoclonal, #H00005863-M02, dilution 1/500). The cells were rinsed with PBS for three times and incubated with the following reagents prepared in PBS supplemented with 1% (w/v) BSA; CF594-conjugated goat anti-mouse IgG1(γ1) (Biotium, #20249), at a final concentration of 2 µg/ml, 4’,6-diamidine-2’-phenylindole dihydrochloride (DAPI) (Merck #10236276001) at a final concentration of 1 µg/ml, and Phalloidin-iFluor 488 Reagent (Abcam, ab176753), 1/100 dilution. The samples were incubated for 40 minutes at room temperature. The samples were finally rinsed with PBS three times, once with milli-Q water and mounted on a slide using Vectashield mounting medium (Vector Laboratories, H-1000-10). Cell images were acqurired using a 2D array scanning laser confocal microscope (Infinity 3, VisiTech) using a 60x/1.4 Plan Apo objective lens (Nikon). Each image comprised 35 serial images with 0.3 µm intervals along the Z-axis to span the full thickness of the cell.

### Quantification of Paxillin structures

In order to appreciate the change in the number of the paxillin structure in a cell, the ratio of the area of paxillin structures and the area of the cytoplasm was quantified using the Fiji Plug-in “Trainable Weka Segmentation” (42,43). Firstly, all 35 Z-slices of each image were stacked and summed to form one image. To this image, “Trainable Weka Segmentation” was applied as stated in the Supporting information S4 and two binary images were generated; one which represents the area of paxillin signals and the other which represents the area of cytoplasm (including the paxillin area). Using these images, the paxillin:cytoplasm ratio was calculated as described in Supporting information S4. For each biological replicate, 20 images were quantified. The data from three biological replicates were analysed with nested t-test using Prism (GraphPad).

## Data availability

The datasets supporting the conclusions of this article are included within the article and its additional file.

## Supporting information

This article contains supporting information provided as Supporting information figures S1 – S6.

## Acknowledgements

We thank Sally Prigent for her helpful advice. The work was conducted using Core Biotechnology Services (CBS) at the University of Leicester; cell imaging was done at the Advanced Imaging Facility, flow cytometry analysis was conducted at the Flow Cytometry Facility, and cell counting and RNA quality checks were done at the Nucleus Genomics (University of Leicester). We thank Reshma R. Vaghela and Nic Sylvius for technical support for the CBS equipment. We thank India-May M. Baker and Dan G. Bowden for their helpful advice on flow cytometry data acquisition and analysis.

## Author contributions

Conceptualisaion: N.L.H., A.M.F., K.H., K.T.; Investigation: N.L.H., L.R.A., S.M.D.A., K.H., KT.; Formal analyses: N.L.H., L.R.A., S.M.D.A. K.R.S., Methodology: N.L.H., K.R.S., K.H., K.T.; Software: K.R.S.,; Writing – original draft: K.T.; Writing – review & editing: N.L.H., L.R.A., K.R.S., A.M.F., K. H.; Supervision: A.M.F. K.T., Project administration: K.T.; Funding acquisition: K.H., K.T.

