## Supporting information Figures S1-S6 for "Endogenous oncogenic KRAS expression increases cell proliferation and motility in near-diploid hTERT RPE-1 cells"

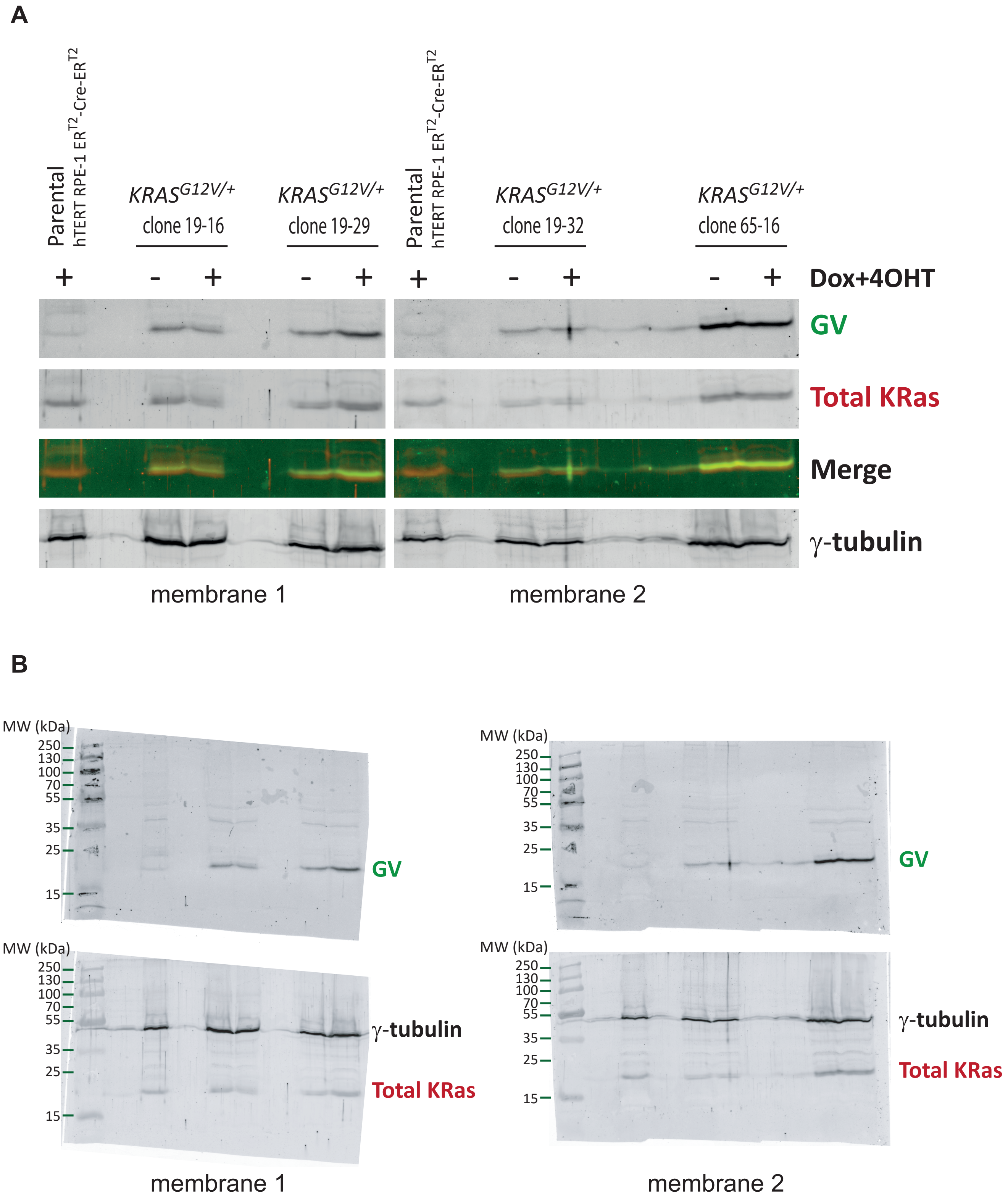
A
Parental
hTERT RPE-1 ER$^{T2}$-Cre-ER$^{T2}$
*KRAS*$^{G12V/+}$ clone 19-16
*KRAS*$^{G12V/+}$ clone 19-29
*KRAS*$^{G12V/+}$ clone 19-32
*KRAS*$^{G12V/+}$ clone 65-16
Dox+4OHT
GV
Total KRas
Merge
γ-tubulin
membrane 1
membrane 2
B
MW (kDa)
250
130
100
70
55
35
25
15
GV
γ-tubulin
Total KRas
membrane 1
membrane 2

Fig. S2A and B

A. The *KRAS* genome structure for two representative products, KRas4A and KRas4B

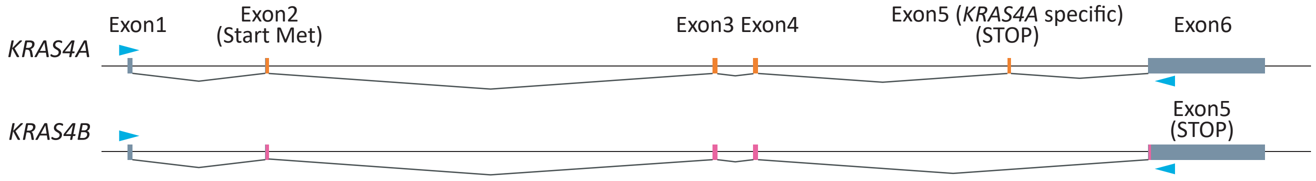


B. *KRAS*$^{G12G(WT)/+}$ cDNA

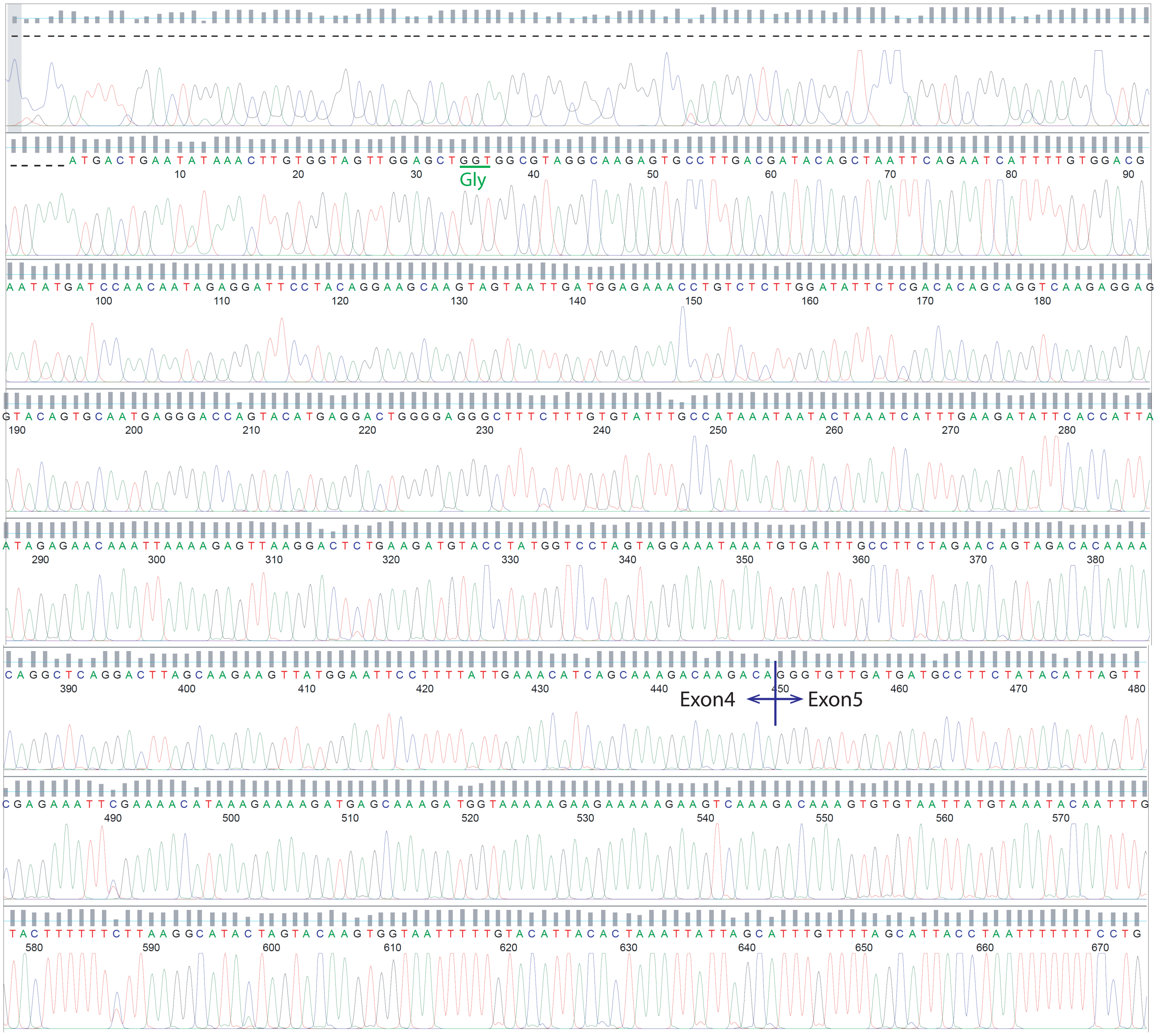

Fig. S2C

C. *KRAS*$^{G12V/+}$ cDNA

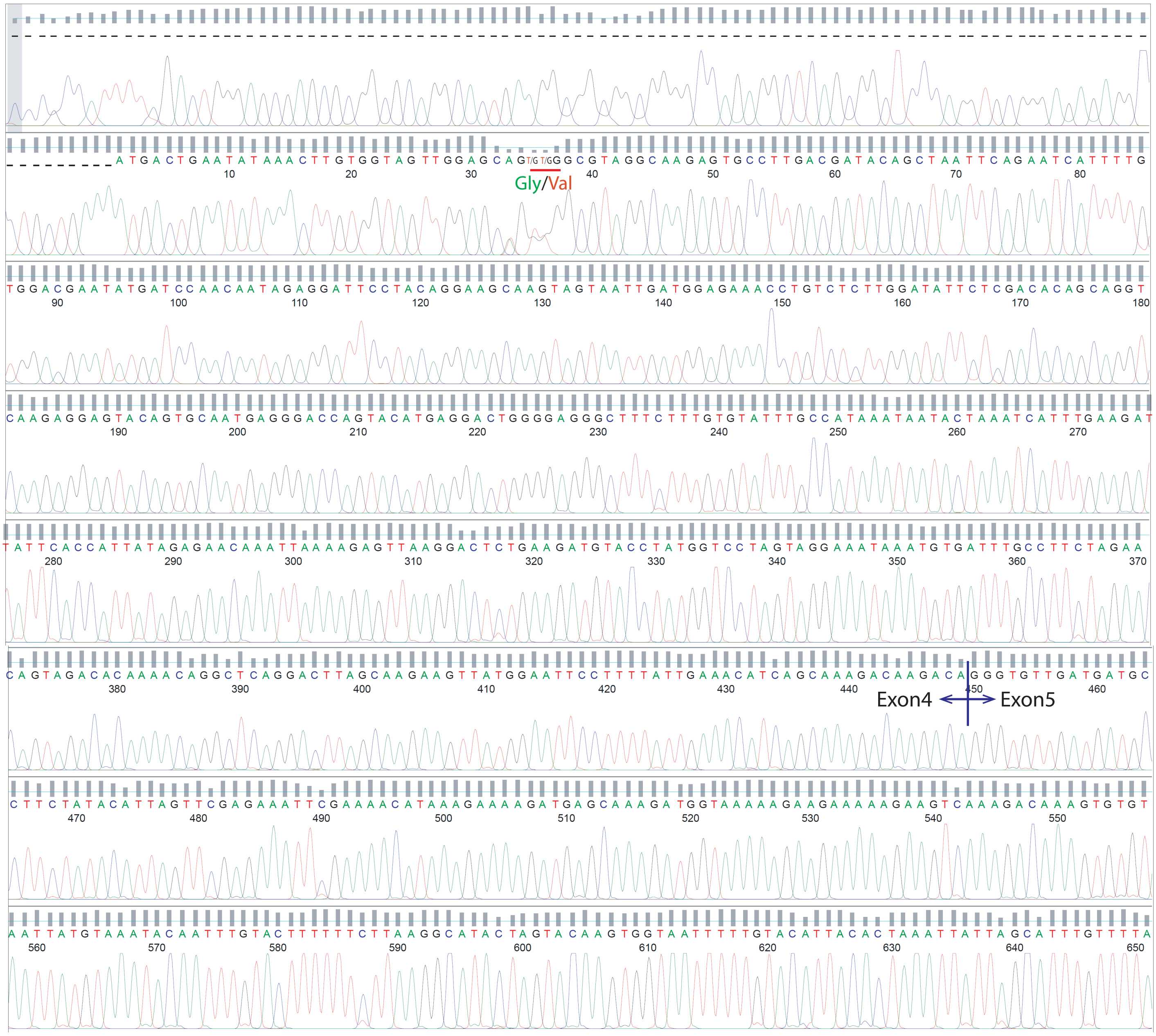

Fig. S2D

D. *KRAS*$^{G12C/+}$ cDNA

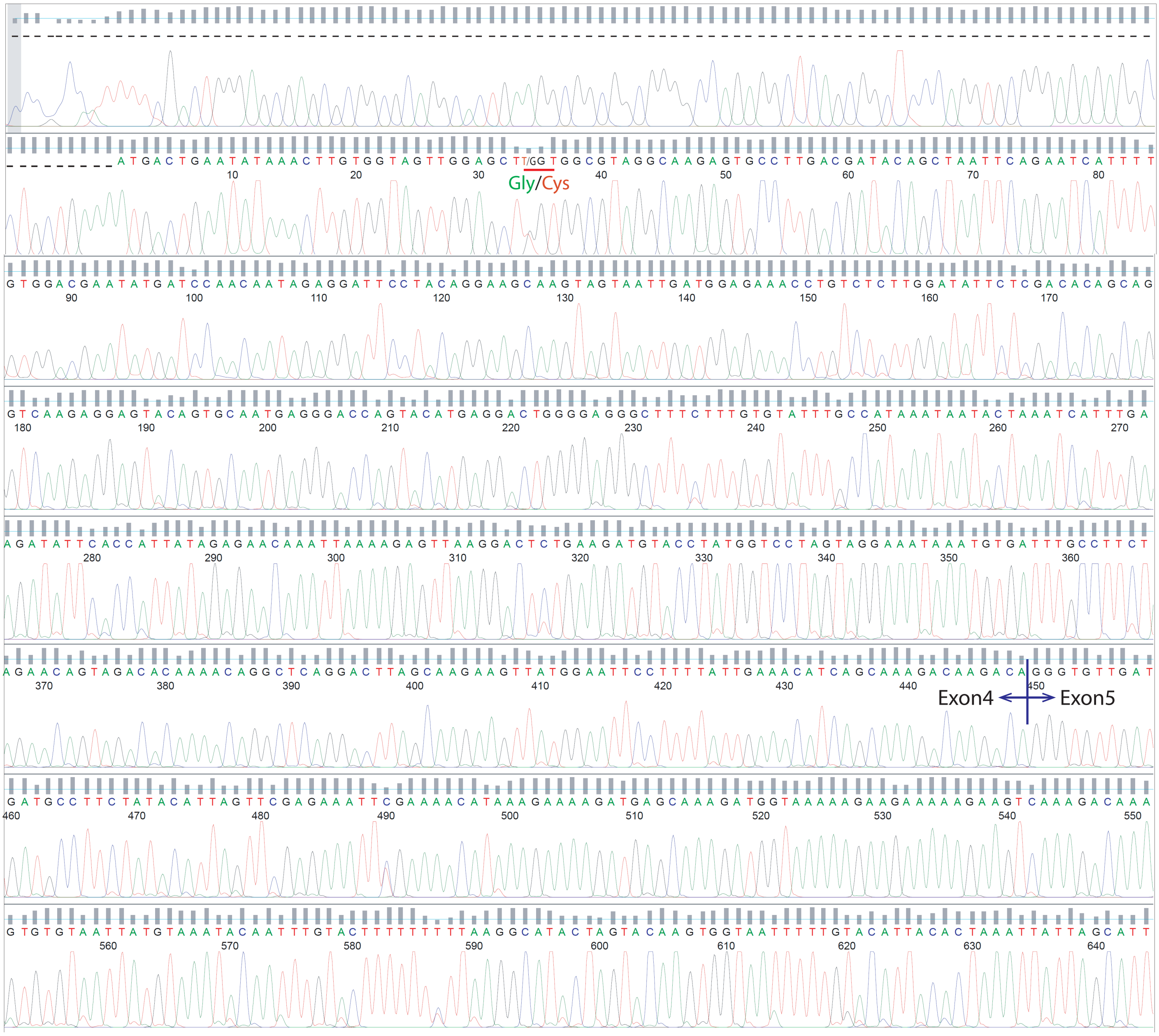

E. *KRAS*$^{G12D/+}$ cDNA

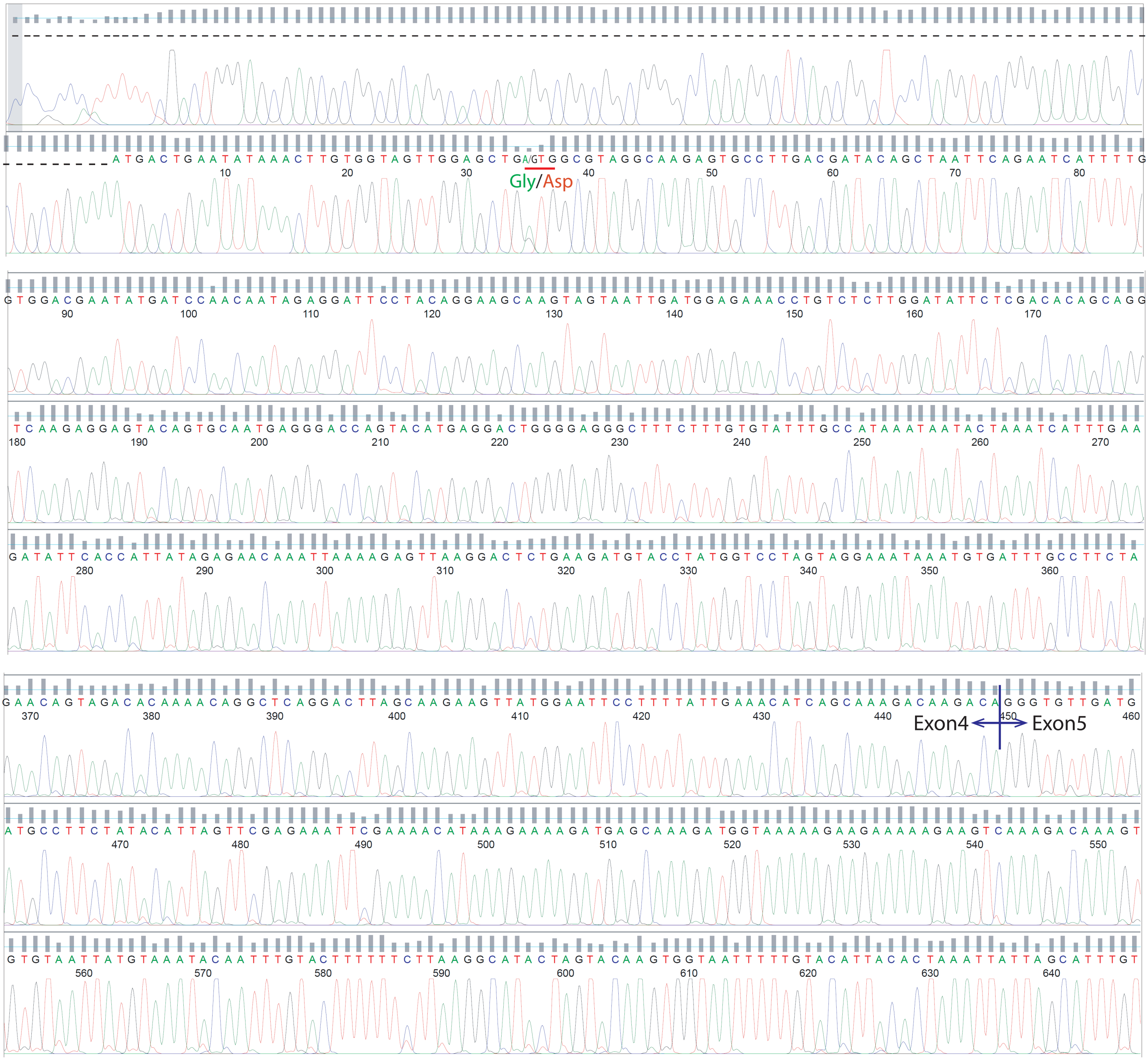

A. $KRAS^{G12G(WT)/+}$ triplicates

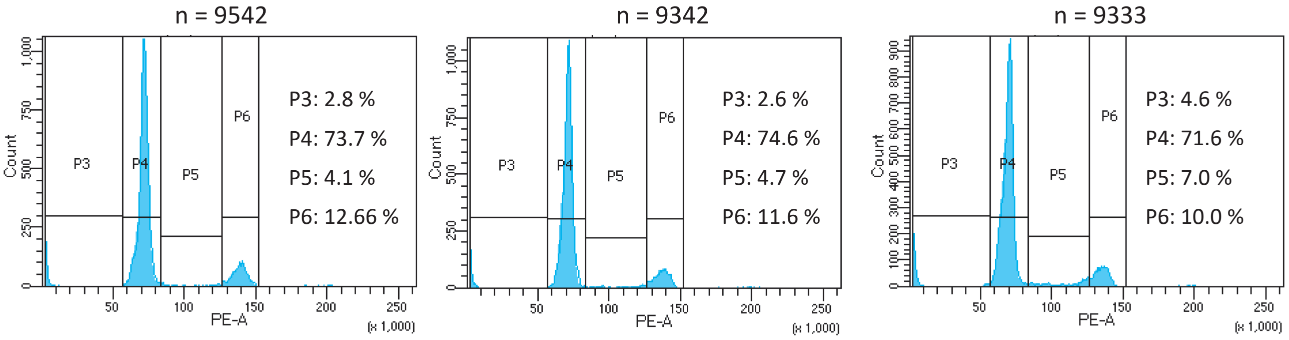


B. $KRAS^{G12V/+}$ triplicates

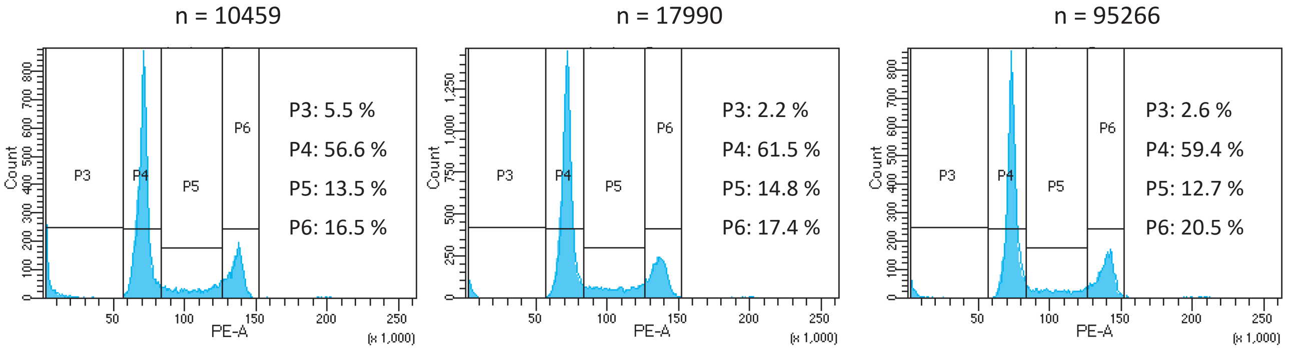

**A**

Trainable Weka Segmentation

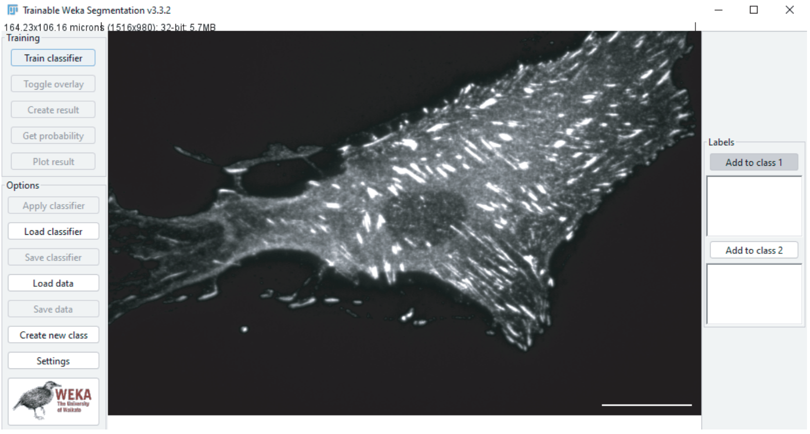


**B**

Segmented Paxillin structure area

Segmented cytoplasmic area

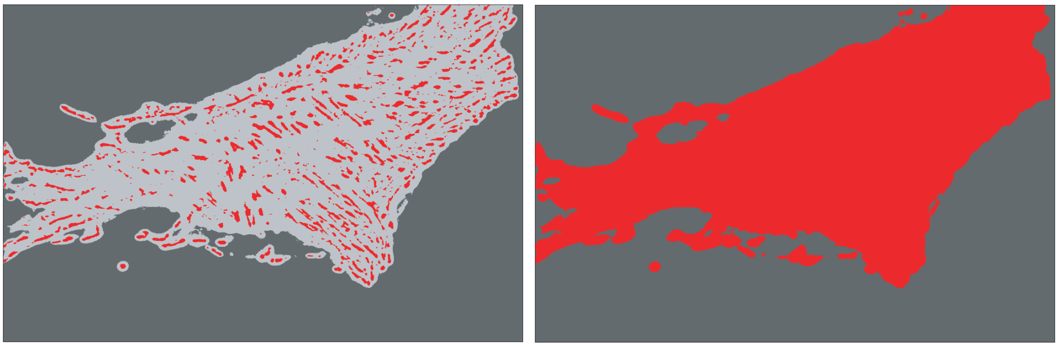

Fig. S5

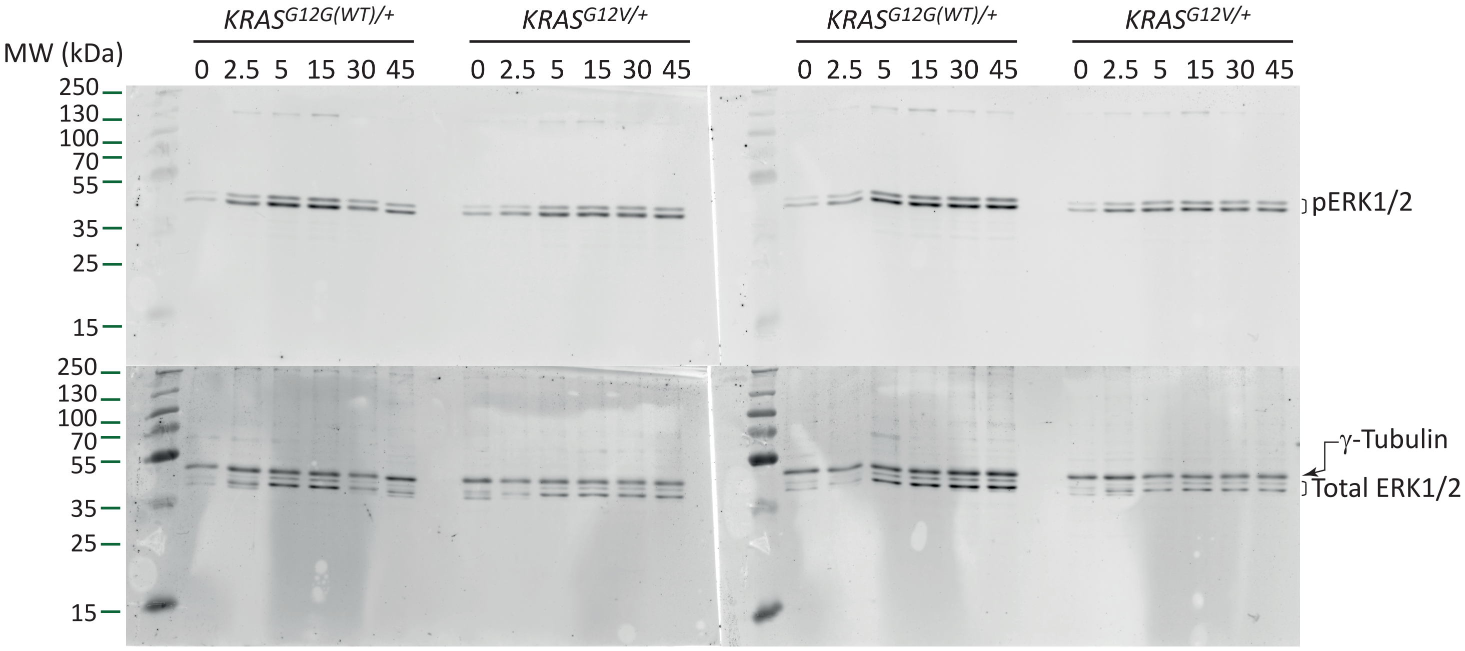


Replicate 1

Replicate 2

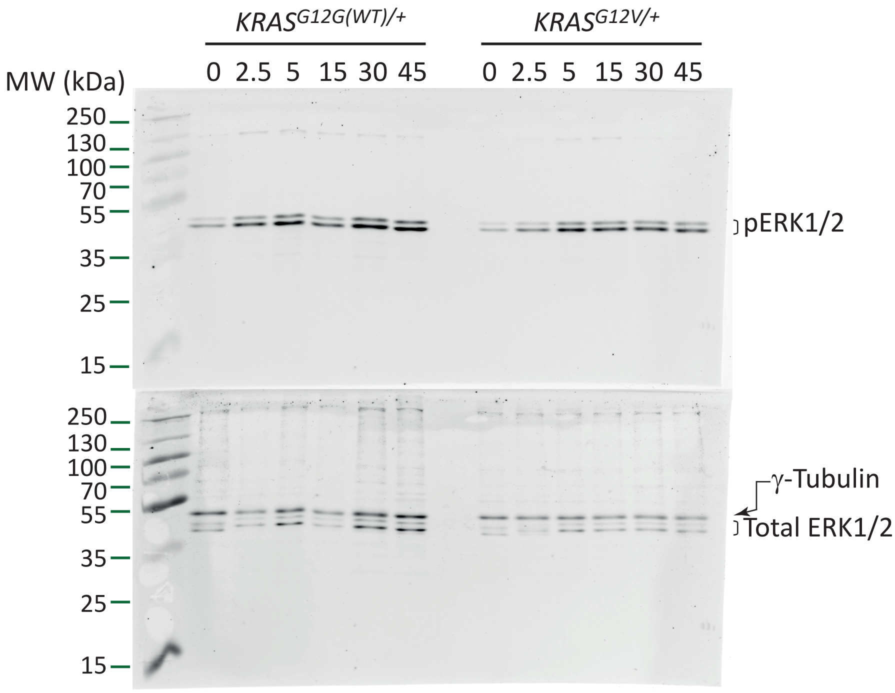


Replicate 3

Fig. S6

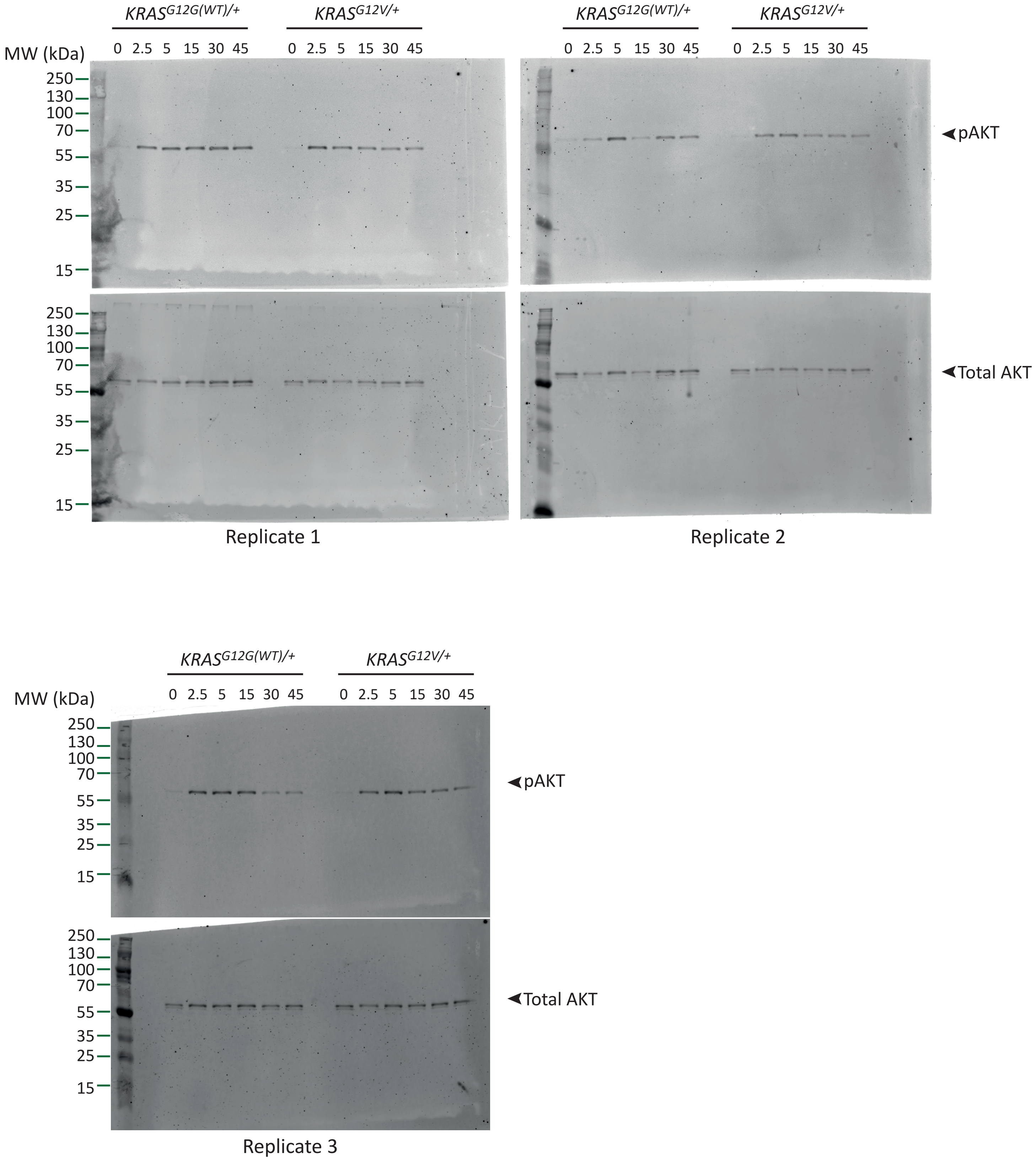

KRAS^G12G(WT)/+
KRAS^G12V/+
0 2.5 5 15 30 45
0 2.5 5 15 30 45
MW (kDa)
250
130
100
70
55
35
25
15
pAKT
Total AKT
Replicate 1
Replicate 2
Replicate 3
